# Natural Killer cells demonstrate distinct eQTL and transcriptome-wide disease associations, highlighting their role in autoimmunity

**DOI:** 10.1101/2021.05.10.443088

**Authors:** James J Gilchrist, Seiko Makino, Vivek Naranbhai, Evelyn Lau, Sara Danielli, Dan Hameiri-Bowen, Wanseon Lee, Esther Ng, Justin Whalley, Julian C Knight, Benjamin P Fairfax

## Abstract

Natural Killer (NK) cells are innate lymphocytes with central roles in immunosurveillance and are implicated in autoimmune pathogenesis. The degree to which regulatory variants affect NK gene expression is poorly understood. We performed expression quantitative trait locus (eQTL) mapping of negatively selected NK cells from a population of healthy Europeans (n=245). We find a significant subset of genes demonstrate eQTL specific to NK cells and these are highly informative of human disease, in particular autoimmunity. An NK cell transcriptome-wide association study (TWAS) across five common autoimmune diseases identified further novel associations at 27 genes. In addition to these *cis* observations, we find novel master-regulatory regions impacting expression of *trans* gene networks at regions including 19q13.4, the Killer cell Immunoglobulin-like Receptor (KIR) Region, *GNLY* and *MC1R*. Our findings provide new insights into the unique biology of NK cells, demonstrating markedly different eQTL from other immune cells, with implications for disease mechanisms.

## Introduction

NK cells are large granular lymphocytes, comprising 5-15% of peripheral blood lymphocytes, and are key innate effector cells[1]. Functions include cytotoxicity towards virally-infected and malignant cells, and production and secretion of cytokines including IFN*γ*, defining NK cells as prototypical Group 1 innate lymphoid cells [2]. NK cells express germline encoded receptors, the best-characterised being Killer cell Immunoglobulin-like Receptor (KIRs), which, dependent on associated intracellular signalling domains, can be activating or inhibitory. Viral infection and dysplasia elicit concomitant induction of stress antigens, triggering activating KIRS, and MHC Class I molecule downregulation, removing the ligands for inhibitory KIRs. Immunodeficiencies characterised by NK cell deficiency[3][4][5][6] or dysfunction[7] are illustrative of the role of NK cells in human health, and are characterised by susceptibility to viral infection, in particular herpesviruses, and early-onset malignancy.

Genome-wide association studies have provided unparalleled insights into the genetic determinants of human health and disease[8]. The majority of associated alleles are non-coding and thought to exert phenotype via the regulation of gene expression. Understanding which cell types and conditions such variants demonstrate activity is vital in determining pathogenic mechanisms. Expression quantitative trait loci (eQTL) analysis of immune cells isolated from blood has demonstrated high degrees of cellular specificity in regulatory variant function. Whilst analyses of NK cell eQTL have been performed, sample sizes have been small and further data are required[9], especially for the identification of *trans* regulatory variants which have smaller effect sizes. We describe an eQTL study performed across 245 healthy individuals of Northern European ancestry. We demonstrate that NK cells display many hundreds of eQTL that are not observed in other cell types, and find evidence of NK cell specific master-regulatory regions. These eQTL colocalise with disease associated loci, providing novel insights into the role of NK cells in human health and disease.

## Methods

### Study samples

Healthy individuals of European ancestry (n=245) were recruited following written informed consent (Oxfordshire Research Ethics Committee COREC reference 06/Q1605/55). Participants had a median age of 28 years (range 18-66) and 117 of 245 were male. PBMCs, purified by Ficoll-Pacque density gradient centrifugation, were isolated from whole blood collected into EDTA-containing tubes (Vacutainer system, Becton Dickinson). Cells were washed in Hanks’ balanced salt solution (Invitrogen) and enumerated with a haemocytometer.

CD56^+^CD3^*-*^ NK cells were isolated by magnetic-activated cell sorting negative selection (MACS, Miltentyi). Cells snap-frozen in RLT reagent (Qiagen) prior to RNA extraction. Total RNA was extracted using RNAeasy Mini kits (Qiagen), quantified using a NanoDrop (ThermoFisher), and BioAnalyzer quantification in a subset.

### Gene expression quantification

Total RNA from 245 individuals was quantified using Illumina HumanHT-12 v4 BeadChip gene expression arrays including 47,231 probes. Probe sequences mapping to more than one genomic locus, and probe sequences containing common genomic variation (minor allele frequency *>* 1%) were excluded from further analysis. Gene expression estimates were normalised (random-spline normalisation) before variance-stabilising transformation using the R package lumi[10]. Samples were processed in 2 batches, and gene expression estimates corrected using the the R package ComBat[11]. Following quality control, gene expression quantified at 29,011 probes, mapping to 18,078 unique genes were included in the analysis.

### Genotyping & imputation

Genomic DNA was extracted from whole blood using Gentra Puregene Blood Kits (Qiagen) according to manufacturer’s instructions, before dsDNA quantification using PicoGreen (Invitrogen). Genome-wide genotypes at 733,202 loci were generated using HumanOmniExpress-12v1.0 BeadChips (Illumina). SNP quality control (QC) filters were applied as follows: minor allele frequency (MAF) *<* 4%, Hardy-Weinberg equilibrium (HWE) *p <* 1 × 10^*-*6^, plate effect *p <* 1 × 10^*-*6^ and SNP missingness*>* 2%. We calculated identity by descent and principal components of LD-pruned genome-wide genotyping data to identify related individuals and sample outliers respectively. Samples with call rates *<* 98% were excluded from analysis. QC metrics were calculated in PLINK.

Following QC, 609,704 variants were taken forward for genome-wide imputation with pre-phasing using SHAPEIT[12] and imputation using IMPUTE2[13]. We used 1000G Phase 1 as a reference panel. Following imputation, SNPs with imputation info scores *<* 0.9 or HWE *p <* 1 × 10^*-*6^ were excluded from further analysis. Genotypes at 6,012,996 loci were taken forward for eQTL mapping. Genotyping QC did not identify population outliers or related samples and all 245 individuals were included in the analysis.

We used SHAPEIT phased genotypes at 100 SNPs in the KIR region on chromosome 19 to perform KIR copy number imputation using KIR*IMP v1.2.0[14] with a UK KIR reference panel. To assess the performance of KIR copy number imputation, we assayed the presence or absence of 12 functional KIR genes (2DL1-5, 2DS1-5, 3DL1, 3DS1) and two pseudogenes (2DP1 and 3DP1) in a subset of the study samples (n=171). For each gene/pseudogene we used sequence-specific primers to amplify PCR products from genomic DNA. PCR products were separated with agarose gel electrophoresis and stained with ethidium bromide. The presence or absence of each KIR gene/pseudogene was confirmed with two complementary PCR reactions, resulting in two PCR products of different length at each locus.

### eQTL analysis

QTLtools[15] was used for eQTL mapping under an additive linear model, including PCs of gene expression data to limit the effect of confounding variation, with *cis* eQTL defined within 1Mb of the associated probe and *trans* loci those *>* 5Mb from associated probe. 32 and 20 PCs maximised eQTL discovery in *cis* and *trans* respectively (Figure S1). Prior to eQTL mapping, expression data was corrected for PCs before rank normal transformation. For *cis* eQTL mapping, QTLtools approximates a permutation test at each phenotype, controlling for multiple testing burden at the level of each phenotype. A second level of multiple-testing correction across all phenotypes tested was applied in R using qvalue[16]. Forward-backward stepwise regression implemented in QTLtools[15] was used to define multiple independent signals reaching a permutation-based significance threshold, mapping lead eSNPs at each signal to define identify secondary, tertiary and quaternary *cis* eQTL. We mapped *trans* eQTL using QTLtools in two stages, firstly fitting additive linear models for each phenotype:genotype pair generating nominal p-values for each association, before determining the significance threshold by permuting phenotypes 100 times. Correlation between phenotypes (but not genotypes) is maintained within each permutation run. For all analyses we considered an *FDR <* 0.05 to be significant.

### Identification of NK cell-specific eQTL

To determine sharing of NK cell *cis* eQTL with other cells, moloc[17] was used in R to compare association at NK cell *cis* eQTL between NK cells and four other previously published datasets of eQTL in primary immune cell subsets from individuals of European ancestry[18][19][20]. Moloc adopts a Bayesian framework to compare models of association across multiple traits at a given genetic locus, using summary statistics. We applied moloc at all identified *cis* eQTL in NK cells, defining the evidence for shared regulatory effects on gene expression across NK cells (this dataset, n=245), CD4^+^ T cells (n=282), CD8^+^ T cells (n=271), monocytes (n=414) and neutrophils (n=101). We considered an eQTL to be unique to NK cells where the posterior probability of the eQTL not being shared with another cell type *>* 0.8, and where a unique eQTL in NK cells was the most likely model overall (either an eQTL in NK cells is the only eQTL at that locus, or where other eQTL at the same locus are distinct from the eQTL in NK cells).

### Allele specific expression

To validate the effect of regulatory variation at *MC1R*, we used the C-BASE allele specific expression assay [21][22], designing primers to full length MC1R and cloning PCR product of cDNA and genomic from NK cells from 5 heterozygous individuals heterozygous using TA Cloning kit (Thermofisher). In addition, the same approach was used from monocyte cDNA samples for a separate 5 heterozygote individuals. Cloning product was transformed and colonies isolated with Taqman genotyping being performed on isolated colonies (spotted with pipette tip into PCR solution) using a probes designed to detect both alleles. 96 colonies were tested for each sample, significance of effect for NK cells was performed using a ChiSquared test (1df), comparing ratios of genomic to cDNA. A paired T test was performed for the monocyte samples.

### Colocalisation & enrichment

We used ENCODE functional annotations[23] to interrogate colocalisation of chromatin states and transcription factor binding sites in lymphoblastoid cell lines (LCLs - GM12878) with eQTL in NK cells. We used ChIP-seq defined transcription factor binding sites for 52 transcription factors, and segmentation of the LCL genome into 7 functional tracks defined by ChromHMM[24] and SegWay[25]; promoter regions, promoter flanking regions, enhancers, weak enhancers or open chromatin cis regulatory elements, CTCF enriched elements, transcribed regions, and repressed or low activity regions. We calculated the density of functional annotations surrounding each *cis* eSNP, counting the number of annotations falling into 1kb bins up to 1Mb from each feature. We used a permutation approach implemented in QTLtools to calculate evidence for enrichment of eSNPs with functional annotations, calculating the frequency of observed overlap between a given functional annotation and an eSNP, comparing this to the the number of overlaps expected by chance (permuting phenotypes across all probes tested). We tested for enriched biological pathways among *cis* NK cell-specific eQTL using the GOBP database in XGR[26]. Genes with *cis* NK cell-specific eQTL were tested against a background of all genes tested in the eQTL analysis, using a hypergeometric test.

We used two approaches to test for colocalisation of causal variants between *cis* eQTL and GWAS traits. Firstly we used Regulatory Trait Concordance (RTC)[27] implemented in QTLtools. RTC integrates GWAS hits, eQTL data and local LD structure to quantify the decrease in eQTL significance when the phenotype is residualised for the GWAS hit. We used RTC to test for colocalisation at *cis* eQTL with 87,919 GWAS-significant loci (*p <* 5 × 10^*-*8^) from 5,823 GWAS studies/traits downloaded on 28/01/2021 from the NHGRI-EBI GWAS Catalog[8] (Table S1). An advantage of the RTC method is that it only requires the identity of GWAS-significant peak SNPs, rather than complete GWAS summary statistics. We considered RTC scores *>* 0.9 to be significant. Secondly, we used the R package coloc[28] to identify evidence of causal variants shared by NK cell eQTL and GWAS loci. Coloc adopts a Bayesian approach to compare evidence for independent or shared association signals for two traits at a given genetic locus. We tested for colocalisation between NK cell primary *cis* eQTL and 100 GWAS traits (Table S2) with evidence of trait-association (*p <* × 10^*-*6^) within a 250kb window of the peak eSNP at each significant NK cell primary *cis* eQTL. We considered a posterior probability *>* 0.8 supporting a shared causal locus to be significant.

To test for enrichment of colocalisation of GWAS traits among NK cell eQTL, we used Fisher Exact tests to compare the proportion of NK cell eQTL within 250kb of a GWAS locus with evidence of trait association (*p <* × 10^*-*6^) for which their is evidence of a shared causal locus, comparing this to the proportion observed for a null GWAS trait (height). Throughout we applied FDR correction to account for the number of annotations and traits tested.

### TWAS analysis

We performed transcriptome-wide association studies (TWAS) implemented in FUSION[29] of five autoimmune diseases, using functional gene expression weights calculated from the NK cell eQTL data described here to impute NK cell gene expression into previously published GWAS of ulcerative colitis[30], Crohn’s disease[30], systemic lupus erythematosus[31], primary biliary cirrhosis[32] and rheumatoid arthritis[33]. For each gene, we correct normalised gene expression for PCs of gene expression (n=32 as for eQTL mapping) and genome-wide genotypes (n=10) and estimate *cis* genetic heritability (SNPs within 1Mb of the TSS), retaining significantly heritable genes (*p <* 0.05). For heritable genes we cross-validated genotype-imputed gene expression estimates, selecting the best-performing model (highest *R*^2^) from 5 prediction models; best linear unbiased predictor, Bayesian linear mixed model, elastic-net regression (*α* = 0.5), LASSO regression and single best eQTL. We then pass functional weights from the best-performing of these models and GWAS summary statistics to FUSION to compute TWAS association statistics for each gene. We applied FDR correction across all genes included in each TWAS, considering *FDR <* 0.05 to be significant. To address whether trait-associated genes were conditionally independent, we performed conditional stepwise regression including all significant TWAS genes on a single chromosome. Finally, to complement the TWAS analysis, at each TWAS-significant gene we used coloc to assess the probability that the GWAS risk locus and NK cell eQTL share a causal variant.

## Results

### *cis* eQTL mapping

NK cells were negatively selected from PBMCs from 245 healthy European adults, and gene expression quantified using Illumina gene expression arrays, giving a readout of 18,078 genes post QC. Whole-genome imputation of genome-wide genotyping data, yielded high-quality genotypes at 6,012,996 autosomal loci. We defined *cis*-acting variants within 1Mb of the TSS, mapping *cis* eQTL under an additive linear model in QTLtools[15]. To alleviate the effects of confounding variation we included PCs of expression data as covariates in the model, finding 32 PCs maximised *cis* eQTL discovery (Figure S1). We identified *cis* eQTL at 3,951 genes of 18,078 tested (21.9%) at *FDR <* 0.05 (Figure 1A, Table S3). By conditioning on the peak eSNP at each significant *cis* eQTL, we further identified 528, 63 and 3 independent eQTLs at secondary, tertiary and quaternary levels respectively (Figure 1A, Table S4).

**Figure 1:**
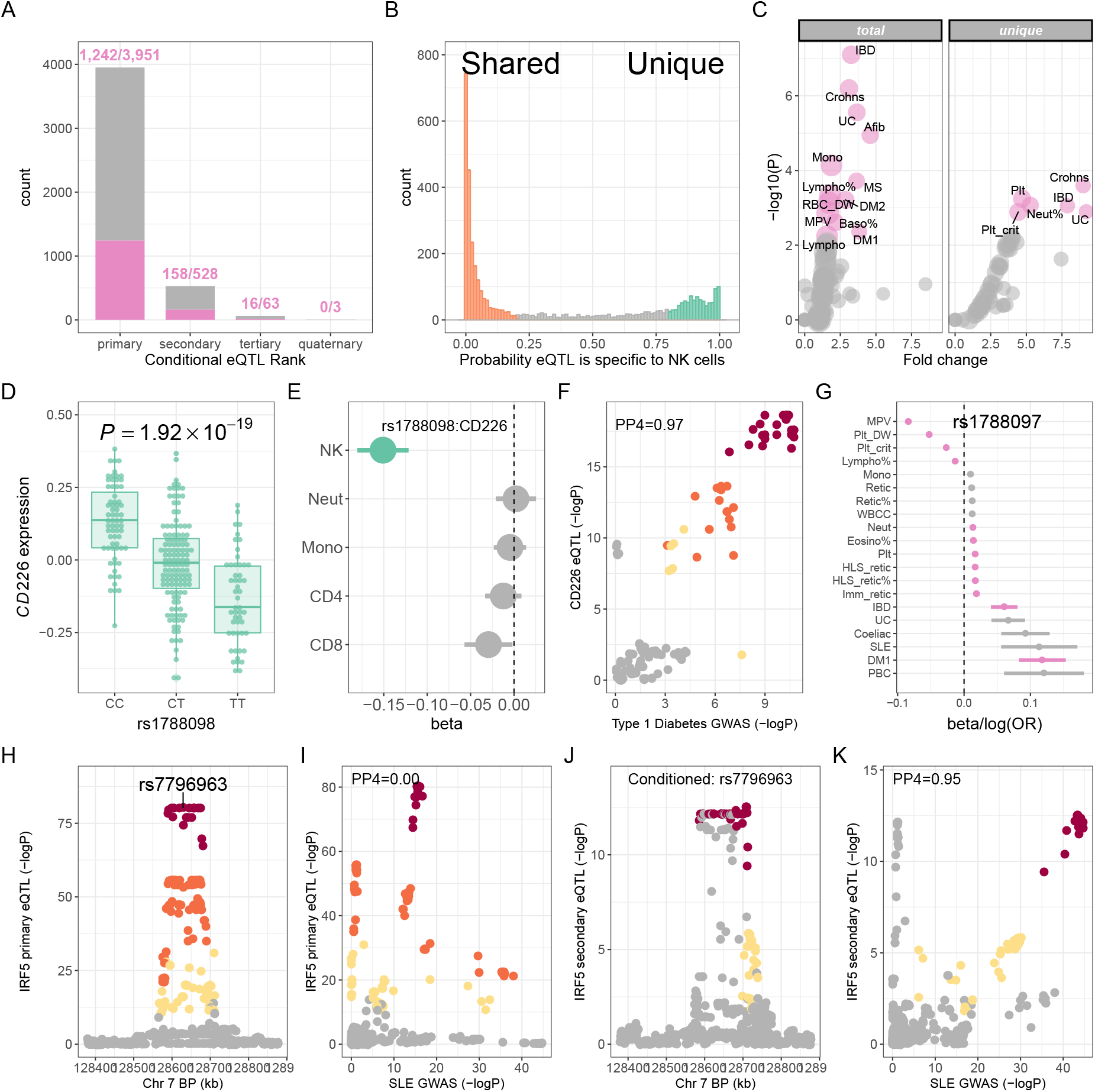
*cis* eQTL in NK cells. (A) Total significant (*FDR <* 0.05) primary and conditional *cis* eQTL in NK cells from 245 individuals. Numbers of eQTL with evidence of colocalisation with at least one GWAS trait (RTC*>* 0.9) are highlighted (pink). (B) Frequencies of *cis* eQTL (*FDR <* 0.05) specific to NK cells or shared with other immune cells; monocytes, neutrophils, CD4^+^ and CD8^+^ T cells. (C) Enrichment of shared causal loci between NK cell eQTL (total, left panel; NK cell-specific, right panel) and GWAS traits (n=100). Traits are compared with the enrichment observed for height as background. Significant traits (*FDR <* 0.05) are highlighted (pink). Point size is proportional to a trait’s number of GWAS-significant loci. (D) Effect of rs1788098 genotype on *CD226* expression in NK cells. (E) The effect of rs1788098 genotype on *CD226* expression is specific to NK cells. Significant eQTL effects are highlighted (green). (F) The *CD226* eQTL in NK cells colocalises with a risk locus for type-1 diabetes. SNPs are coloured according to strength of LD (CEU population) to the peak eSNP (rs1788098); brown *r*^2^ *>* 0.8, orange 0.5 *< r*^2^ ≤ 0.8, yellow 0.2 *< r*^2^ ≤ 0.5, grey *r*^2^ ≤ 0.2. (G) Association of rs1788097 (exact proxy for rs1788098 in European populations, *r*^2^ = 1) with autoimmune diseases and haematological indices. GWAS-significant (*p <* 5 × 10^*-*8^) associations are highlighted (pink). (H) Regional association plot of the primary *IRF5* eQTL in NK cells. (I) The primary *IRF5* eQTL does not colocalise with a GWAS risk locus in the *IRF5* region for systemic lupus erythematosus. (J) Conditionning on the peak *IRF5* eSNP reveals an independent, secondary eQTL for *IRF5* in NK cells. (K) The secondary *IRF5* eQTL does colocalise with the *IRF5* region for systemic lupus erythematosus.

We used a Bayesian approach, implemented in moloc[17], to compare the the evidence for shared or independent effects at NK eQTL (this dataset, n=245) with CD4^+^ T cells[20] (n=282), CD8^+^ T cells[20] (n=271), monocytes[18] (n=414) and neutrophils[19] (n=101). Whilst we find evidence (*PP >* 0.8) supporting sharing of a single regulatory effect at 2,161/3,951 (54.7%) NK eQTL with at least one other cell type, the data best-supports (*PP >* 0.8 model of association being an eQTL in NK cells alone or independent to eQTLs in other cell types) observed NK eQTL being unique to NK cells at 588/3,951 (14.9%) genes (Figure 1B, Table S5). Amongst these NK cell-specific eQTL, 270/588 (49.1%) regulate expression of genes for which there are other independent eQTL in at least one other immune cell. Thus, whilst many NK-cell specific eQTL may be reflective of cell-restricted expression, half represent differential regulation of widely expressed genes.

### *cis* eQTL function

Regulatory genetic variation acting in *cis* operates, at least in part, through *cis*-acting regulatory elements, with eQTL significantly enriched within regions of active chromatin, promoters, enhancers and TSS[34]. To assess the distribution of *cis* eQTLs in NK cells we plotted densities of observed eQTL around functional chromatin states and transcription factor binding sites, calculating relative enrichment of NK eQTL around each feature (Figure S2, Table S6). NK *cis* eQTLs demonstrate clustering around TSS (*p <* 2 × 10^*-*8^), transcribed regions (*p <* 2 × 10^*-*8^), enhancers (*p* = 0.0015) and weak enhancers (*p* = 0.0087), and are underrepresented in regions of the genome with repressed activity (*p <* 2 × 10^*-*8^). Cell-type specific eQTL and conditional eQTL have been demonstrated to show greater overlap with distal regulatory elements, as compared to primary, shared eQTL[35]. In keeping with this, secondary conditional eQTL in NK cells are enriched for enhancers (*p* = 0.00043), but show no enrichment around promoters (Figure S2). Similarly, while NK cell-specific eQTL demonstrate clustering at promoter sites (*p <* 2.2 × 10^*-*7^) and no significant clustering around enhancers, they demonstrate enrichment for weak enhancers (*p* = 0.00053), significantly more so than NK eQTL shared across immune cells, (*p* = 0.0019, fold change = 3.93).

We similarly observed significant enrichment of overlap between NK cell eQTL and transcription factor binding sites, demonstrating enrichment at 44/52 (84.6%) of transcription factors tested (Figure S2, Table S6). The transcription factor binding site enrichment observed across NK cell eQTL encompasses a broad range of transcription factors, many of which are active in immune cells (e.g. NFKB, IRF7, PAX5). In particular, ETS1 has been previously demonstrated to direct NK cell differentiation[36]. We observed no significant enrichment between secondary conditional eQTL in NK cells and transcription factor binding sites. Among NK cell-specific eQTL, we identified significant enrichment of overlap at 14/52 (26.9%) of transcription factor binding sites, including EGR1, POU2F2 and ZEB1 which are predicted to act as transcription factor repressors active in determining ILC subset plasticity[37].

To better understand the biological pathways subject to regulatory variation specific to NK cells, we performed enrichment analysis of genes with an NK cell-specific eQTL (against a background of all tested genes) in XGR[26]. In that analysis, NK cell-specific eQTL are enriched for genes within 26 biological pathways annotated by Gene Ontology Biological Processes (GOBP, Table S7, Figure S3). Enriched biological processes highlight the established biology of NK cells, including pathways mediating cell death (GO:0006915, apoptotic process; GO:1902042, negative regulation of extrinsic apoptotic signalling pathway via death domain receptors; GO:0097190, apoptotic signalling pathway; GO:0042981, regulation of apoptotic process; GO:0006919, activation of cysteine-type endopeptidase activity involved in apoptotic process), anti-viral host defence (GO:0009615, response to virus; GO:0051607, defense response to virus), and innate immune signalling (GO:0032496, response to lipopolysaccharide; GO:0033209, tumour necrosis factor-mediated signalling pathway)

### *cis* eQTL disease associations

To investigate the disease informativeness of NK cells we adopted two complementary approaches. We firstly used RTC to determine the proportion of NK cell eQTL predicted to share a causal variant with NHGRI-EBI GWAS Catalog studies (n=5,823). Among primary *cis* NK cell eQTL, 1,242/3,951 (31.4%) eQTL are predicted to share the same functional variant with at least one GWAS trait (Figure 1A). Among eQTL predicted to be unique to NK cells, 161/588 (27.4%) loci are predicted to colocalise with at least one GWAS-significant locus. Conditional eQTL are similarly informative of human disease and GWAS traits, with 158/528 (29.9%), 16/63 (25.3%) and 0/3 secondary, tertiary and quaternary conditional eQTL in NK cells predicted to colocalise with GWAS trait-associated loci.

Next, to better understand the impact that regulatory variation modifying gene expression in NK cells has on human health and disease, we used evidence of colocalisation between NK cell eQTL and GWAS traits, derived with coloc[28], to estimate the degree of enrichment of NK eQTL colocalisation with GWAS traits as compared to that expected by chance. Height GWAS loci are enriched for regulatory variation operating in connective tissue and mesenchymal stem cells but not immune cells[38], and any shared causal variants with NK cell eQTL are likely to be representative of the proportion one would expect by chance (i.e. shared causal loci are likely to represent effects mediated by other cell types)[39]. We therefore used a well-powered GWAS of height in European populations (UK Biobank, URL: http://www.nealelab.is/uk-biobank/) as our background against which to compare enrichment in other traits. We performed enrichment analysis across 100 traits (Table S2); 40 UK Biobank continuous traits (anthropometrics and haematological indices) and 60 NHGRI-EBI GWAS Catalog traits (well-powered, case-control studies in populations of European ancestry). In that analysis, we identify 13 GWAS traits with significant enrichment (FDR*<* 0.05) of colocalisation with NK cell eQTL, and 6 traits with significant enrichment of colocalisation with NK cell-specific eQTL (Figure 1C, Table S2). Among both NK cell-specific and shared eQTL, we observe marked enrichment of shared causal loci with autoimmune diseases (inflammatory bowel disease, multiple sclerosis, type-1 diabetes) and a range of haematological indices including monocyte and lymphocyte count (Figure 1C). NK cell-specific eQTL are enriched for colocalisation with platelet indices (platelet count and crit).

Genes with NK cell-specific eQTL include *CD226* (Figure 1D-E), which encodes CD226 (also known as DNAX accessory molecule-1), a surface-expressed immunoglobulin superfamily glycoprotein, widely expressed on NK cells [40]. CD226 engages CD155 on antigen-presenting cells, an interaction increasing cytotoxicity and IFN*γ* production, that is competitively inhibited by NK-expressed TIGIT and CD96[41]. The NK cell-specific eQTL for *CD226* (peak eSNP: rs1788098, *p* = 1.92 × 10^*-*19^) colocalises with genetic loci for a range of haematological indices and autoimmune diseases; including IBD and T1DM (Figure 1F-G). The direction of effect of *CD226* expression on autoimmune disease risk is of reduced expression increasing disease risk, suggesting less robust NK cell responses (potentially directed at viral pathogens or autoreactive T cells) increases the risk of autoimmunity. Indeed, NK cells have been shown to restrain the activity of CD4^+^ T cells in a CD226-dependent manner in the context of multiple sclerosis[42]. Other NK cell-specific eQTL informative for human disease include; the caspase protease *CASP8* (peak eSNP: rs3769821, *p* = 1.24 × 10^*-*8^) which colocalises with risk loci for multiple malignancies (breast cancer, non-small cell lung cancer, melanoma, oesophageal squamous cell carcinoma), and the fucosyltransferase *FUT11* (peak eSNP: rs11000765, *p* = 1.91 × 10^*-*18^) which colocalises with asthma and lung function.

As expected, NK eQTL shared with other immune cells are also highly informative with respect to human health and disease. For instance, an *ERAP2* eQTL (peak eSNP: rs1363974, *p* = 1.33 × 10^*-*76^), predicted to be shared across all immune cells tested (NK cells, CD4^+^ and CD8^+^ T cells, monocytes and neutrophils), is a genetic determinant of neutrophil and lymphocyte percentage (Figure S4). As described above, conditional eQTL are also informative with regard to human disease risk (Table S4). A secondary eQTL for *IRF5*, encoding the transcription factor IRF5, a positive regulator of type-1 interferon production, provides an example of this (Figure 1H-K). Here the primary eQTL at *IRF5* (peak eSNP: rs7796963, *p* = 2.37 × 10^*-*65^) has no evidence of colocalisation with human disease traits, however the secondary eQTL (peak eSNP: rs17424921, *p* = 5.39 × 10^*-*9^) shows strong evidence of a shared causal variant with systemic lupus erythematosus.

### *trans* eQTL

Detecting *trans* eQTL within purified cell types can elucidate cell-specific regulatory networks[18]. We defined *trans*-acting regulatory variation as variants *>* 5Mb from the TSS, and tested for associations at 18,078 genes under an additive linear in QTLtools[15], alleviating confounding variation through the inclusion of expression data PCs as covariates in the model (20 PCs maximising *trans* discovery - Figure S1). We identified 5,783 SNPs demonstrating *trans* effects (*FDR <* 0.05) to 116 genes (Figure 2A, Table S8). Much of *trans*-acting regulatory variation is secondary to *cis*-effects on upstream regulatory genes[**westra2013systematic**][43]. Consistent with this, of the 116 genes with *trans* eQTL, 21 have evidence of *cis*-mediation (colocalisation probability of *cis* and *trans* effect *>* 0.8) from 15 genes (Table S8).

**Figure 2:**
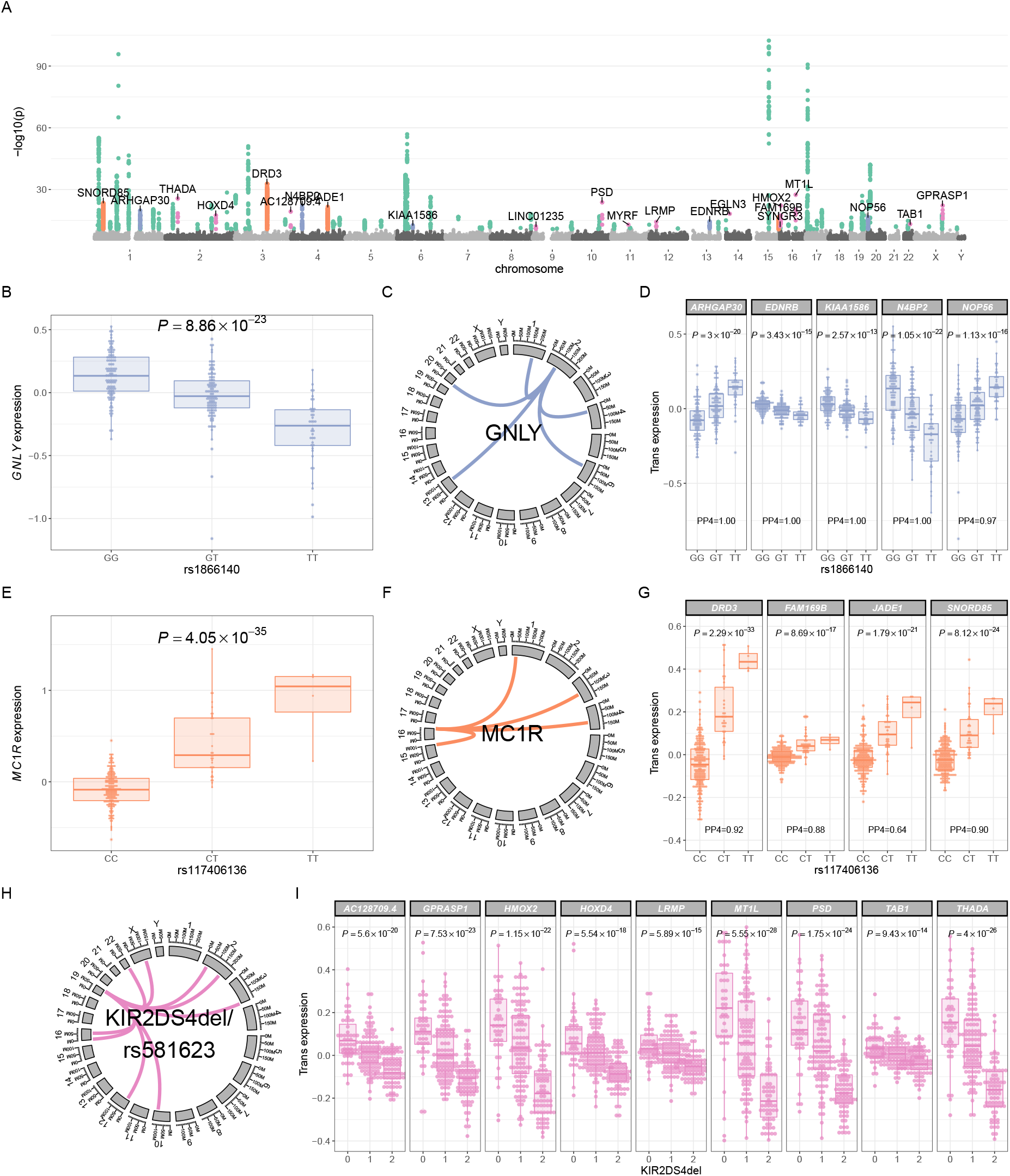
*trans* eQTL in NK cells. (A) Manhattan plot depicting significant *trans* eQTL in NK cells. Physical position (x axis) represents location of the target gene. Coloured points represent significant (*FDR <* 0.05) *trans* eQTL; 5,783 *trans* SNP-gene associations, affecting expression of 116 genes in 87 *trans* regulatory networks. Regulatory networks are highlighted as follows; *GNLY* (blue), *MC1R* (orange), KIR (pink), all others (green). (B) Effect of rs1866140 genotype on *GNLY* expression in NK cells. (C) A *trans* regulatory network of 5 genes, mediated in *cis* by *GNLY* expression. (D) Effect of rs1866140 genotype on *GNLY* regulatory network genes in *trans*. PP4, posterior probability of a shared causal variant with *GNLY cis* eQTL. (E) Effect of rs117406136 genotype on *MC1R* expression in NK cells. (F) A *trans* regulatory network of 4 genes, mediated in *cis* by *MC1R* expression. (G) Effect of rs117406136 genotype on *MC1R* regulatory network genes in *trans*. PP4, posterior probability of a shared causal variant with *MC1R cis* eQTL. (H) A *trans* regulatory network of 9 genes, regulated by rs581623 genotype (proxy for KIR2DS4del). (I) Effect of KIR2DS4del copy number on KIR region regulatory network genes in *trans*. (J) Regional association plot depicting effect of KIR region genotype and KIR copy number on *MT1L* expression in *trans*. SNPs are coloured grey, KIR types coloured green. Significant associations are highlighted (pink).

Among *trans*-acting loci, 8 *trans* eSNPs affect more than one gene distally, and show master-regulatory properties. For instance, a *trans* regulatory network mediated by a NK cell-specific *cis* eQTL for *GNLY* (peak eSNP: rs1866140, *p* = 8.86 × 10^*-*23^), determines expression of 5 genes in *trans*; *ARHGAP30, EDNRB, KIAA1586, N4BP2, NOP56* (Figure 2B-D). *GNLY* encodes granulysin, a secreted cytolytic molecule with broad antimicrobial activity and cytotoxicity towards infected and malignant cells[44]. In addition to granulysin’s cytolytic activity, granulysin also induces immune cell chemotaxis and proinflammatory cytokine production in monocytes. The expression of a secreted molecule as a *cis*-mediator of a *trans* regulatory network within a cell type is highly analogous to the previously-described *trans* eQTL network mediated by lysozyme expression[45][46][47], and suggests a model in which *trans* networks can be mediated by signalling between immune cells, with the afferent component here provided by secreted granulysin.

A second example of a *trans* regulatory network, affecting *DRD3, FAM169B, JADE1* and *SNORD85* expression, is mediated in *cis* by an eSNP affecting *MC1R* expression (Figure 2E-G, peak eSNP: rs117406136, *p* = 4.05 × 10^*-*35^). The *MC1R cis* eQTL is shared with CD4^+^ and CD8^+^ T cells (Table S5), and the lead SNP in the non-imputed genotyping was rs2228479, encoding a non-synonymous polymorphism (V92M) within *MC1R* and in perfect LD (R^2^ = 1) with rs117406136. Given this, we tested for allele specific expression of this allele across five individuals heterozygous for rs2228479, using the C-BASE assay [21][22], observing for all individuals significantly increased expression of the minor allele (*p* = 0.01-*p* = 2.3 × 10^*-*7^, combined *p <* 2.2 × 10^*-*16^, *χ*^2^ test). Notably no ASE was observed in monocytes at this SNP (Figure S5). *MC1R* encodes the melanocortin-1 receptor (MC1R), a high affinity receptor for *α*-melanocyte stimulating hormone (*α*MSH), and most robustly expressed in melanocytes, with associations with tanning[48], freckling, hair and eye colour[49], melanoma[50] and non-melanoma skin cancer[51]. As well as its effects on pigmentation, MC1R/*α*MSH signalling has well-established immunomodulatory functions, and MCR1 is surface expressed on B cells, T cells and NK cells[52]. Whilst the effect of MC1R/*α*MSH signalling in NK cells is undefined, NK cell dysfunction, as part of broader immunodeficiency, is well-characterised as part of syndromic disorders of hypopigmentation, e.g. Chediak-Higashi, Hermansky-Pudlak and Griscelli syndromes[53]. All of these syndromes are characterised by impaired secretory lysosome function dysfunction, impairing melanin secretion from melanocytes as well as cytotoxic granule secretion from NK cells and cytotoxic T cells. Our description here of an NK cell *trans* regulatory network mediated by MC1R expression suggests that the NK cell immune defect seen in syndromic disorders of hypopigmentation may not be confined to defective cytotoxic degranulation.

The largest *trans* regulatory network we observed is a gene network for which the master regulator maps to the leukocyte receptor complex (LRC, 19q13.4), which encodes KIR genes. Expression of 9 genes (*FDR <* 0.05) in *trans* to the LRC are regulated by a KIR region SNP: rs581623 (Figure 2H-I). The LRC is highly polymorphic in terms of KIR gene content and allelic variation. To better define the *cis* mediator of this regulatory network, we imputed KIR copy number in the study samples using KIR*IMP[14] and remapped *trans* eQTL in NK cells using KIR copy number imputations. KIR gene/pseudogene copy numbers were well-imputed (Table S9), with estimated imputation accuracies ranging from 76.2% (KIR2DS2) to 96.9% (KIR2DS1) and high levels of concordance with gene presence/absence as determined by PCR (80.7-100%). In that analysis, rs581623 genotype is an exact proxy for KIR2DS4del: a 22 base-pair deletion in *KIR2DS4* which introduces a premature stop codon[54]. KIR haplotypes are broadly grouped into A and B haplotypes, and for A haplotypes KIR2DS4 is the only activating receptor. KIR2DS4 ligands have been challenging to identify but include HLA-C*05:01-bound peptides derived from epitopes common to a number of bacterial pathogens[55]. KIR2DS4del/rs581623 is the lead regulatory variant for all 9 genes within the KIR *trans* regulatory network (Figure 2I, Figure 2J, Table S10), with no signifiant independent associations remaining after conditioning on KIR2DS4del copy number (Figure S5). In addition, KIR imputation revealed a second KIR *trans* regulatory network independent of KIR2DS4del (Figure S6). That network comprises 3 genes (*MYRF, EGLN3, SYNGR3*) regulated by *KIR3DP1* copy number.

### TWAS

A striking feature of our eQTL analysis was the degree of enrichment for autoimmune disease that we observed among *cis* eQTL in NK cells. To further investigate this we utilised NK cell eQTL data and GWAS summary statistics for autoimmune disease to identify novel autoimmune risk genes within five autoimmune TWAS; ulcerative colitis[30], rheumatoid arthritis[33], systemic lupus erythematosus[31], primary biliary cirrhosis[32] and Crohn’s disease[30]. We identified 98 TWAS-significant, independent gene-trait associations (*FDR <* 0.05) for autoimmune diseases (Figure 3, Supplementary Tables 11-15); ulcerative colitis 21/2,998 genes tested, rheumatoid arthritis 5/3,015 genes tested, systemic lupus erythematosus 18/3,045 genes tested, primary biliary cirrhosis 18/3,035 genes tested, Crohn’s disease 31/3,077 genes tested. We considered a gene-trait association to be novel if there was no GWAS-significant locus within 1Mb of the gene in the trait’s GWAS used as input for the TWAS, and if there was no colocalisation evidence (L2G score ≥ 0.3) supporting gene-trait association in Open Targets Genetics[56] (URL: https://genetics.opentargets.org, accessed 28/01/2021). Of the 98 TWAS-significant trait-associated genes, 27 define novel gene-trait associations (Table 1).

**Table 1:**
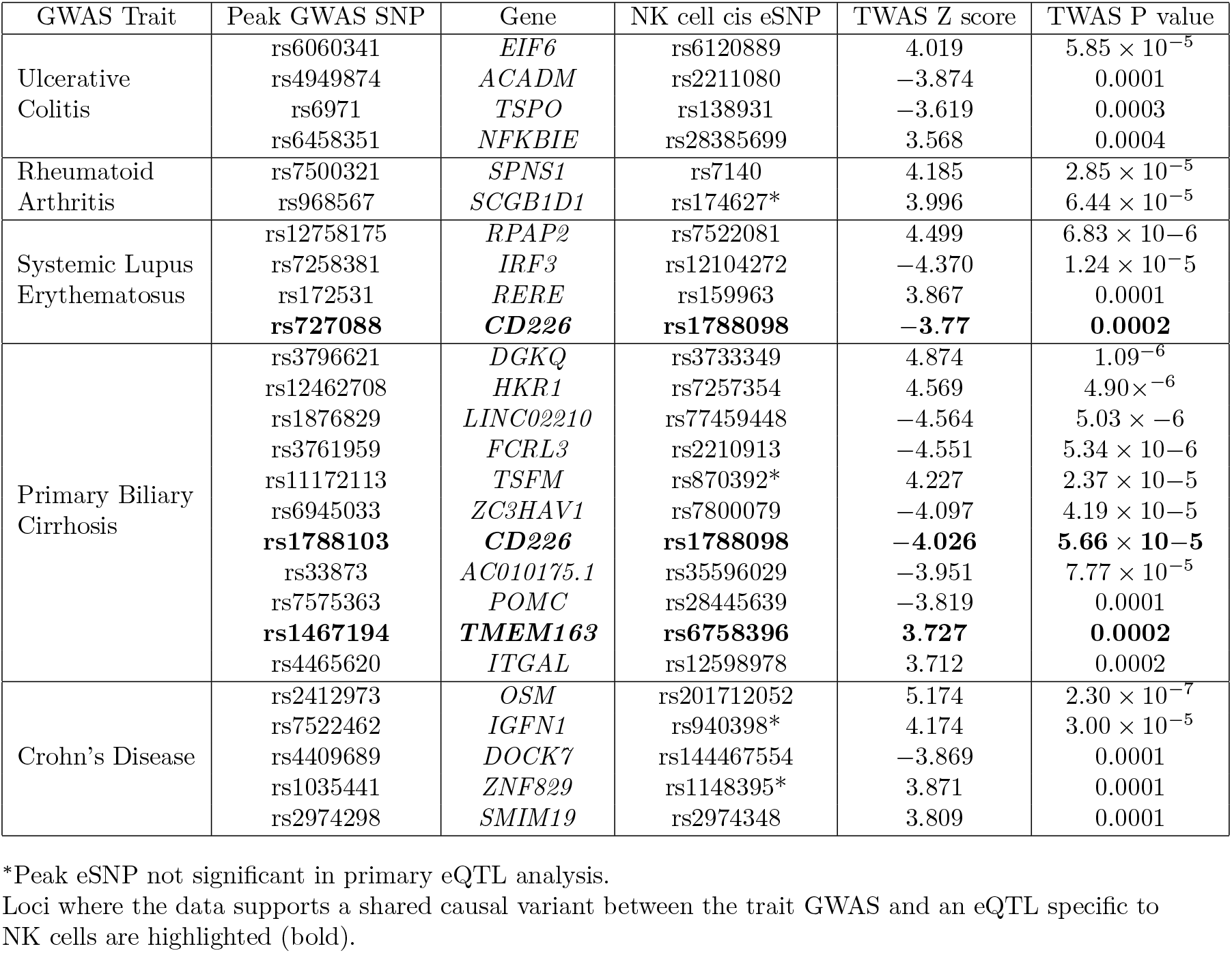
Novel gene-trait associations identified by TWAS.

**Figure 3:**
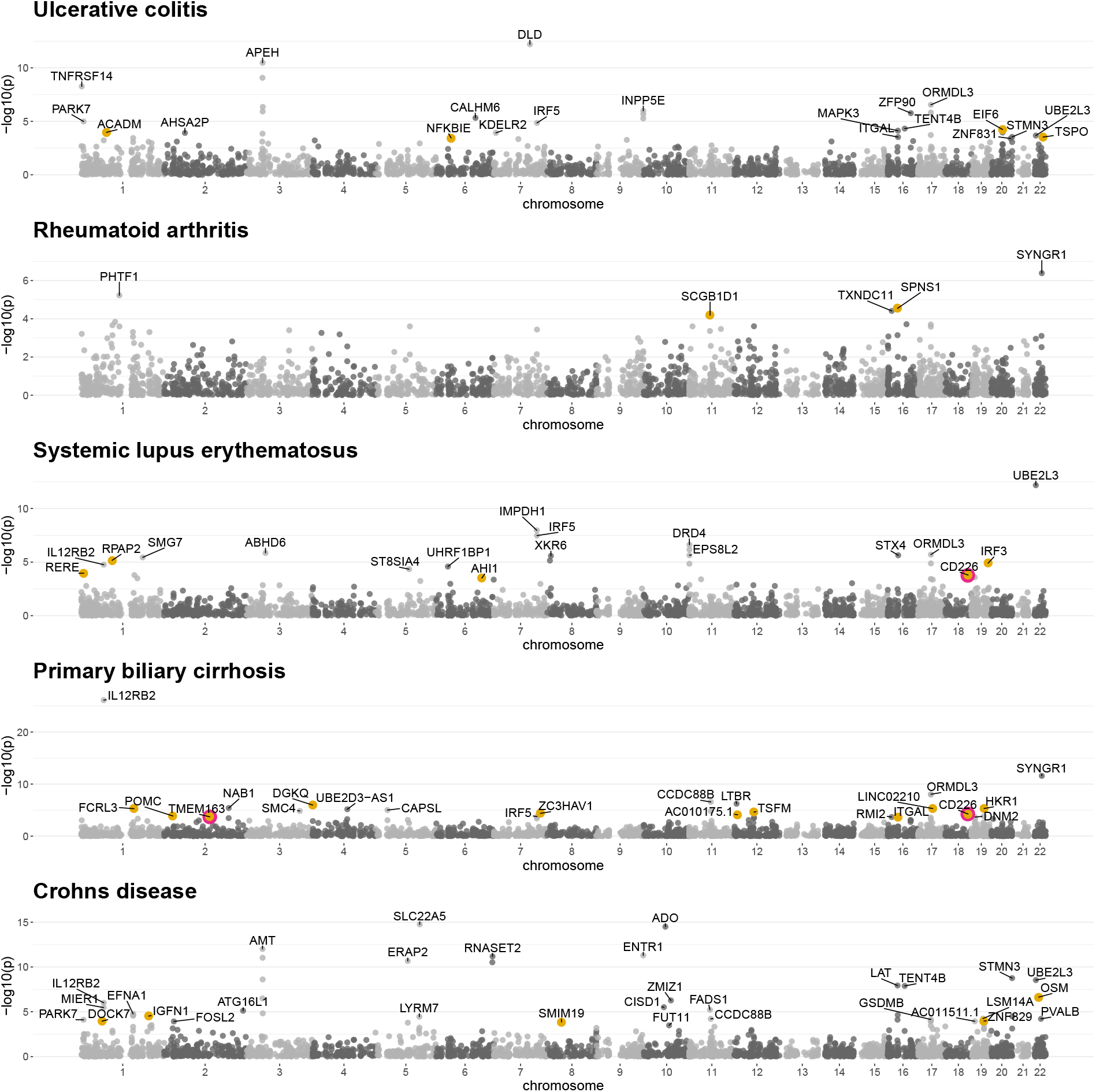
NK cell TWAS in autoimmune diseases. Manhattan plot of autoimmune disease TWAS. All significant trait-associated genes (*n* = 98) are labelled. Novel gene-trait associations (no GWAS-significant locus within 1Mb and no evidence supporting gene-trait association in Open Targets Genetics) are highlighted (yellow, *n* = 27). Novel gene trait associations which colocalise (posterior probability colocalisation *>* 0.8) with an NK cell-specific eQTL are circled (pink, *n* = 3).

We were particularly interested in identifying novel gene-trait associations for which the evidence supports a shared causal variant between the trait GWAS and an eQTL specific to NK cells (defining colocalisation and NK cell specificity as above). We were able to identify three instances of this. Complementary to our identification of NK cell expression *CD226* as a risk factor for type-1 diabetes and multiple sclerosis (Figure 1), we identify NK cell expression *CD226* as a determinant of both systemic lupus erythematosus and primary biliary cirrhosis. We also identified NK cell expression of *TMEM163* as a determinant of primary biliary cirrhosis. *TMEM163* encodes a transmembrane zinc transporter[57], which is hypothesised to mediate zinc accumulation into lysosomes[58]. Notably, zinc deficiency has been reported to be associated with reduced NK cell cytolytic activity[59]. Genetic variation affecting *TMEM163* expression has not previously been implicated in the pathogenesis of autoimmune disease. However, in an interesting parallel to our description of a *trans* regulatory network in NK cells mediated by *MC1R* expression, eQTL for *TMEM163* in whole blood[60] colocalise (PPH4=0.96, Open Targets Genetics) with genetic determinants of hair colour (UK Biobank, URL: http://www.nealelab.is/uk-biobank/). *TMEM163* is not regulated in *trans* by the *MC1R* locus in NK cells, however these data suggest a model in which the shared biological determinants of pigmentation and inflammation operate in NK cells and modify risk of immune-mediated disease.

## Discussion

Understanding the genetic determinants of gene expression in immune cells has proved highly informative in advancing our understanding of immune-mediated disease. Here, we have defined the regulatory landscape of gene expression in primary human NK cells. In doing so, we identified thousands of eQTL operating in NK cells, and demonstrated that a significant proportion of this regulatory variation appears specific to these cells.

Regulatory variation active in NK cells is highly enriched for disease-associated genetic variation. By combining our eQTL mapping with colocalisation analysis, incorporating the results of multiple genetic association studies, we elucidate many examples where regulatory variation specific to NK cells is associated with human disease risk. As such, our data directly implicate gene expression in NK cells in a range of human traits and diseases, most strikingly for autoimmune diseases. We expand on these findings, leveraging our NK eQTL data to perform TWAS in five autoimmune diseases, defining novel roles for NK cell expression of *CD226* and *TMEM163* in systemic lupus erythematosus and primary biliary cirrhosis.

Our findings are in keeping with an emerging role for NK cells in autoimmune disease. This role is supported by observational data across a number of autoimmune diseases, identifying decreased NK cell number in the periphery as a marker of disease risk or activity[61][62][63], and accumulation of NK cells at sites of autoimmunity[64, 65]. The presence of NK cells at sites of autoimmunity could suggest a model in which NK cells directly contribute to tissue damage through cytolysis and/or cytokine secretion, or alternatively a model in which NK cells have an immunoregulatory role, restraining the activity of autoreactive immune cells. Recent data have highlighted the importance of the latter model, defining immunoregulatory roles for NK cells in controlling the activity of CD4^+^ T cells[42][66].

We identified multiple *trans* eQTL operating in NK cells, most notably *trans* regulatory networks mediated by MC1R, granulysin and the KIR region. The identification of a large *trans* regulatory network mediated by a common deletion in an activating KIR receptor (KIR2DS4del) is particularly intriguing and is analogous to previous descriptions of master-regulatory variation to the MHC[67][46]. Variation in KIR type has been implicated in the pathogenesis of infectious and autoimmune disease. It is highly likely that the KIR-mediated *trans* regulatory network we have identified here will have important roles in human disease risk, and that this role will emerge as the KIR region is better-defined, through imputation[14] or sequencing[68] approaches, in GWAS cohorts.

In summary, we have described the *cis* and *trans* regulatory landscapes of gene expression in human NK cells. We have integrated these data with epigenetic and GWAS datasets deriving important insights into NK cell biology and its impact on human health and disease.

## Acknowledgements

J.C.K. is supported by Wellcome Trust Investigator Award [204969/Z/16/Z], NIHR Oxford Biomedical Research Centre and Chinese Academy of Medical Sciences (CAMS) Innovation 537 Fund for Medical Science (grant number: 2018-I2M-2-002), Wellcome Trust Grants 090532/Z/09/Z and 203141/Z/16/Z to core facilities Wellcome Centre for Human Genetics, Oxford Biomedical Research Computing (BMRC) facility, a joint development between the Wellcome Centre for Human Genetics and the Big Data Institute supported by Health Data Research UK and the NIHR Oxford Biomedical Research Centre. Study was funded by Wellcome Trust Intermediate Clinical Fellowship to B.P.F. (no. 201488/Z/16/Z). J.J.G. is funded by a National Institute for Health Research (NIHR) Clinical Lectureship. The views expressed are those of the author(s) and not necessarily those of the NHS, the NIHR or the Department of Health and Social Care.

## References

1. Campbell, K. S. & Hasegawa, J. Natural killer cell biology: an update and future directions. eng. J Allergy Clin Immunol 132, 536–544. ISSN: 1097-6825 (Electronic); 0091-6749 (Print); 0091-6749 (Linking) (Sept. 2013).

2. Spits, H. et al. Innate lymphoid cells —a proposal for uniform nomenclature. Nature Reviews Immunology 13, 145–149. https://doi.org/10.1038/nri3365 (2013).

3. Biron, C. A., Byron, K. S. & Sullivan, J. L. Severe herpesvirus infections in an adolescent without natural killer cells. eng. N Engl J Med 320, 1731–1735. ISSN: 0028-4793 (Print); 0028-4793 (Linking) (June 1989).

4. Gineau, L. et al. Partial MCM4 deficiency in patients with growth retardation, adrenal insufficiency, and natural killer cell deficiency. eng. J Clin Invest 122, 821–832. ISSN: 1558-8238 (Electronic); 0021-9738 (Print); 0021-9738 (Linking) (Mar. 2012).

5. Hanna, S., Béziat, V., Jouanguy, E., Casanova, J. L. & Etzioni, A. A homozygous mutation of RTEL1 in a child presenting with an apparently isolated natural killer cell deficiency. eng. J Allergy Clin Immunol 136, 1113–1114. ISSN: 1097-6825 (Electronic); 0091-6749 (Linking) (Oct. 2015).

6. Cottineau, J. et al. Inherited GINS1 deficiency underlies growth retardation along with neutropenia and NK cell deficiency. eng. J Clin Invest 127, 1991–2006. ISSN: 1558-8238 (Electronic); 0021-9738 (Print); 0021-9738 (Linking) (May 2017).

7. Grier, J. T. et al. Human immunodeficiency-causing mutation defines CD16 in spontaneous NK cell cytotoxicity. eng. J Clin Invest 122, 3769–3780. ISSN: 1558-8238 (Electronic); 0021-9738 (Print); 0021-9738 (Linking) (Oct. 2012).

8. Buniello, A. et al. The NHGRI-EBI GWAS Catalog of published genome-wide association studies, targeted arrays and summary statistics 2019. eng. Nucleic Acids Res 47, D1005–D1012. ISSN: 1362-4962 (Electronic); 0305-1048 (Print); 0305-1048 (Linking) (Jan. 2019).

9. Schmiedel, B. J. et al. Impact of Genetic Polymorphisms on Human Immune Cell Gene Expression. Cell 175, 1701–1715.e16 (Nov. 2018).

10. Du, P., Kibbe, W. A. & Lin, S. M. lumi: a pipeline for processing Illumina microarray. eng. Bioinformatics 24, 1547–1548. ISSN: 1367-4811 (Electronic); 1367-4803 (Linking) (July 2008).

11. Johnson, W. E., Li, C. & Rabinovic, A. Adjusting batch effects in microarray expression data using empirical Bayes methods. eng. Biostatistics 8, 118–127. ISSN: 1465-4644 (Print); 1465-4644 (Linking) (Jan. 2007).

12. Delaneau, O., Marchini, J. & Zagury, J.-F. A linear complexity phasing method for thousands of genomes. eng. Nat Methods 9, 179–181. ISSN: 1548-7105 (Electronic); 1548-7091 (Linking) (Dec. 2011).

13. Howie, B. N., Donnelly, P. & Marchini, J. A flexible and accurate genotype imputation method for the next generation of genome-wide association studies. eng. PLoS Genet 5, e1000529. ISSN: 1553-7404 (Electronic); 1553-7390 (Print); 1553-7390 (Linking) (June 2009).

14. Vukcevic, D. et al. Imputation of KIR Types from SNP Variation Data. The American Journal of Human Genetics 97, 593–607. ISSN: 0002-9297. http://www.sciencedirect.com/science/article/pii/S0002929715003699 (2015).

15. Delaneau, O. et al. A complete tool set for molecular QTL discovery and analysis. Nature Communications 8, 15452. https://doi.org/10.1038/ncomms15452 (2017).

16. Storey, J. D. A direct approach to false discovery rates. Journal of the Royal Statistical Society: Series B (Statistical Methodology) 64, 479–498. eprint: https://rss.onlinelibrary.wiley.com/doi/pdf/10.1111/1467-9868.00346. https://rss.onlinelibrary.wiley.com/doi/abs/10.1111/1467-9868.00346 (2002).

17. Giambartolomei, C. et al. A Bayesian framework for multiple trait colocalization from summary association statistics. eng. Bioinformatics 34, 2538–2545. ISSN: 1367-4811 (Electronic); 1367-4803 (Print); 1367-4803 (Linking) (Aug. 2018).

18. Fairfax, B. P. et al. Innate immune activity conditions the effect of regulatory variants upon monocyte gene expression. eng. Science 343, 1246949. ISSN: 1095-9203 (Electronic); 0036-8075 (Print); 0036-8075 (Linking) (Mar. 2014).

19. Naranbhai, V. et al. Genomic modulators of gene expression in human neutrophils. eng. Nat Commun 6, 7545. ISSN: 2041-1723 (Electronic); 2041-1723 (Linking) (July 2015).

20. Kasela, S. et al. Pathogenic implications for autoimmune mechanisms derived by comparative eQTL analysis of CD4+ versus CD8+ T cells. eng. PLoS Genet 13, e1006643. ISSN: 1553-7404 (Electronic); 1553-7390 (Print); 1553-7390 (Linking) (Mar. 2017).

21. Rainbow, D. B. et al. Commonality in the genetic control of Type 1 diabetes in humans and NOD mice: variants of genes in the IL-2 pathway are associated with autoimmune diabetes in both species. Biochem Soc Trans 36, 312–5 (June 2008).

22. Heap, G. A. et al. Genome-wide analysis of allelic expression imbalance in human primary cells by high-throughput transcriptome resequencing. Hum Mol Genet 19, 122–34 (Jan. 2010).

23. Dunham, I. et al. An integrated encyclopedia of DNA elements in the human genome. Nature 489, 57–74. https://doi.org/10.1038/nature11247 (2012).

24. Ernst, J. & Kellis, M. ChromHMM: automating chromatin-state discovery and characterization. Nature Methods 9, 215–216. https://doi.org/10.1038/nmeth.1906 (2012).

25. Hoffman, M. M. et al. Unsupervised pattern discovery in human chromatin structure through genomic segmentation. Nature Methods 9, 473–476. https://doi.org/10.1038/nmeth.1937 (2012).

26. Fang, H., Knezevic, B., Burnham, K. L. & Knight, J. C. XGR software for enhanced interpretation of genomic summary data, illustrated by application to immunological traits. Genome Medicine 8, 129. https://doi.org/10.1186/s13073-016-0384-y (2016).

27. Nica, A. C. et al. Candidate causal regulatory effects by integration of expression QTLs with complex trait genetic associations. eng. PLoS Genet 6, e1000895. ISSN: 1553-7404 (Electronic); 1553-7390 (Print); 1553-7390 (Linking) (Apr. 2010).

28. Giambartolomei, C. et al. Bayesian test for colocalisation between pairs of genetic association studies using summary statistics. eng. PLoS Genet 10, e1004383. ISSN: 1553-7404 (Electronic); 1553-7390 (Print); 1553-7390 (Linking) (May 2014).

29. Gusev, A. et al. Integrative approaches for large-scale transcriptome-wide association studies. eng. Nat Genet 48, 245–252. ISSN: 1546-1718 (Electronic); 1061-4036 (Print); 1061-4036 (Linking) (Mar. 2016).

30. Jostins, L. et al. Host–microbe interactions have shaped the genetic architecture of inflammatory bowel disease. Nature 491, 119–124. https://doi.org/10.1038/nature11582 (2012).

31. Bentham, J. et al. Genetic association analyses implicate aberrant regulation of innate and adaptive immunity genes in the pathogenesis of systemic lupus erythematosus. eng. Nat Genet 47, 1457– 1464. ISSN: 1546-1718 (Electronic); 1061-4036 (Print); 1061-4036 (Linking) (Dec. 2015).

32. Cordell, H. J. et al. International genome-wide meta-analysis identifies new primary biliary cirrhosis risk loci and targetable pathogenic pathways. eng. Nat Commun 6, 8019. ISSN: 2041-1723 (Electronic); 2041-1723 (Linking) (Sept. 2015).

33. Okada, Y. et al. Genetics of rheumatoid arthritis contributes to biology and drug discovery. Nature 506, 376–381. https://doi.org/10.1038/nature12873 (2014).

34. Gaffney, D. J. et al. Dissecting the regulatory architecture of gene expression QTLs. eng. Genome Biol 13, R7. ISSN: 1474-760X (Electronic); 1465-6906 (Print); 1474-7596 (Linking) (Jan. 2012).

35. Dobbyn, A. et al. Landscape of Conditional eQTL in Dorsolateral Prefrontal Cortex and Colocalization with Schizophrenia GWAS. eng. Am J Hum Genet 102, 1169–1184. ISSN: 1537-6605 (Electronic); 0002-9297 (Print); 0002-9297 (Linking) (June 2018).

36. Taveirne, S. et al. The transcription factor ETS1 is an important regulator of human NK cell development and terminal differentiation. Blood 136, 288–298. https://doi.org/10.1182/blood.2020005204 (Jan. 2020).

37. Pokrovskii, M. et al. Characterization of Transcriptional Regulatory Networks that Promote and Restrict Identities and Functions of Intestinal Innate Lymphoid Cells. eng. Immunity 51, 185–197. ISSN: 1097-4180 (Electronic); 1074-7613 (Print); 1074-7613 (Linking) (July 2019).

38. Wood, A. R. et al. Defining the role of common variation in the genomic and biological architecture of adult human height. Nature Genetics 46, 1173–1186. https://doi.org/10.1038/ng.3097 (2014).

39. Zhernakova, D. V. et al. Identification of context-dependent expression quantitative trait loci in whole blood. eng. Nat Genet 49, 139–145. ISSN: 1546-1718 (Electronic); 1061-4036 (Linking) (Jan. 2017).

40. Shibuya, A. et al. DNAM-1, a novel adhesion molecule involved in the cytolytic function of T lymphocytes. eng. Immunity 4, 573–581. ISSN: 1074-7613 (Print); 1074-7613 (Linking) (June 1996).

41. Martinet, L. & Smyth, M. J. Balancing natural killer cell activation through paired receptors. Nature Reviews Immunology 15, 243–254. https://doi.org/10.1038/nri3799 (2015).

42. Gross, C. C. et al. Impaired NK-mediated regulation of T-cell activity in multiple sclerosis is reconstituted by IL-2 receptor modulation. eng. Proc Natl Acad Sci U S A 113, E2973–82. ISSN: 1091-6490 (Electronic); 0027-8424 (Print); 0027-8424 (Linking) (May 2016).

43. Battle, A. et al. Characterizing the genetic basis of transcriptome diversity through RNA-sequencing of 922 individuals. eng. Genome Res 24, 14–24. ISSN: 1549-5469 (Electronic); 1088-9051 (Print); 1088-9051 (Linking) (Jan. 2014).

44. Sparrow, E. & Bodman-Smith, M. D. Granulysin: The attractive side of a natural born killer. eng. Immunol Lett 217, 126–132. ISSN: 1879-0542 (Electronic); 0165-2478 (Linking) (Jan. 2020).

45. Rotival, M. et al. Integrating genome-wide genetic variations and monocyte expression data reveals trans-regulated gene modules in humans. PLoS Genet 7, e1002367 (Dec. 2011).

46. Fairfax, B. P. et al. Genetics of gene expression in primary immune cells identifies cell type-specific master regulators and roles of HLA alleles. eng. Nat Genet 44, 502–510. ISSN: 1546-1718 (Electronic); 1061-4036 (Print); 1061-4036 (Linking) (Mar. 2012).

47. Ruffieux, H. et al. EPISPOT: an epigenome-driven approach for detecting and interpreting hotspots in molecular QTL studies. bioRxiv (2020).

48. Visconti, A. et al. Genome-wide association study in 176,678 Europeans reveals genetic loci for tanning response to sun exposure. Nature Communications 9, 1684. https://doi.org/10.1038/s41467-018-04086-y (2018).

49. Eriksson, N. et al. Web-based, participant-driven studies yield novel genetic associations for common traits. eng. PLoS Genet 6, e1000993. ISSN: 1553-7404 (Electronic); 1553-7390 (Print); 1553-7390 (Linking) (June 2010).

50. Law, M. H. et al. Genome-wide meta-analysis identifies five new susceptibility loci for cutaneous malignant melanoma. eng. Nat Genet 47, 987–995. ISSN: 1546-1718 (Electronic); 1061-4036 (Print); 1061-4036 (Linking) (Sept. 2015).

51. Chahal, H. S. et al. Genome-wide association study identifies novel susceptibility loci for cutaneous squamous cell carcinoma. eng. Nat Commun 7, 12048. ISSN: 2041-1723 (Electronic); 2041-1723 (Linking) (July 2016).

52. Neumann Andersen, G. et al. MC(1) receptors are constitutively expressed on leucocyte subpopulations with antigen presenting and cytotoxic functions. eng. Clin Exp Immunol 126, 441–446. ISSN: 0009-9104 (Print); 1365-2249 (Electronic); 0009-9104 (Linking) (Dec. 2001).

53. Dotta, L. et al. Clinical, laboratory and molecular signs of immunodeficiency in patients with partial oculo-cutaneous albinism. eng. Orphanet J Rare Dis 8, 168. ISSN: 1750-1172 (Electronic); 1750-1172 (Linking) (Oct. 2013).

54. Maxwell, L. D., Wallace, A., Middleton, D. & Curran, M. D. A common KIR2DS4 deletion variant in the human that predicts a soluble KIR molecule analogous to the KIR1D molecule observed in the rhesus monkey. eng. Tissue Antigens 60, 254–258. ISSN: 0001-2815 (Print); 0001-2815 (Linking) (Sept. 2002).

55. Sim, M. J. W. et al. Human NK cell receptor KIR2DS4 detects a conserved bacterial epitope presented by HLA-C. Proceedings of the National Academy of Sciences 116, 12964 (June 2019).

56. Ghoussaini, M. et al. Open Targets Genetics: systematic identification of trait-associated genes using large-scale genetics and functional genomics. Nucleic Acids Research 49, D1311–D1320. https://doi.org/10.1093/nar/gkaa840 (Feb. 2021).

57. Sanchez, V. B., Ali, S., Escobar, A. & Cuajungco, M. P. Transmembrane 163 (TMEM163) protein effluxes zinc. eng. Arch Biochem Biophys 677, 108166. ISSN: 1096-0384 (Electronic); 0003-9861 (Print); 0003-9861 (Linking) (Nov. 2019).

58. Cuajungco, M. P. & Kiselyov, K. The mucolipin-1 (TRPML1) ion channel, transmembrane-163 (TMEM163) protein, and lysosomal zinc handling. eng. Front Biosci (Landmark Ed) 22, 1330– 1343. ISSN: 1093-4715 (Electronic); 1093-4715 (Linking) (Mar. 2017).

59. Allen, J. I., Perri, R. T., McClain, C. J. & Kay, N. E. Alterations in human natural killer cell activity and monocyte cytotoxicity induced by zinc deficiency. eng. J Lab Clin Med 102, 577–589. ISSN: 0022-2143 (Print); 0022-2143 (Linking) (Oct. 1983).

60. Vósa, U. et al. Unraveling the polygenic architecture of complex traits using blood eQTL metaanalysis. bioRxiv, 447367 (Jan. 2018).

61. Bossowski, A., Urban, M. & Stasiak-Barmuta, A. Analysis of circulating T gamma/delta lymphocytes and CD16/56 cell populations in children and adolescents with Graves’ disease. eng. Pediatr Res 54, 425–429. ISSN: 0031-3998 (Print); 0031-3998 (Linking) (Sept. 2003).

62. Cameron, A. L., Kirby, B. & Griffiths, C. E. M. Circulating natural killer cells in psoriasis. eng. Br J Dermatol 149, 160–164. ISSN: 0007-0963 (Print); 0007-0963 (Linking) (July 2003).

63. Park, Y.-W. et al. Impaired differentiation and cytotoxicity of natural killer cells in systemic lupus erythematosus. eng. Arthritis Rheum 60, 1753–1763. ISSN: 0004-3591 (Print); 0004-3591 (Linking) (June 2009).

64. Dotta, F. et al. Coxsackie B4 virus infection of beta cells and natural killer cell insulitis in recent-onset type 1 diabetic patients. eng. Proc Natl Acad Sci U S A 104, 5115–5120. ISSN: 0027-8424 (Print); 1091-6490 (Electronic); 0027-8424 (Linking) (Mar. 2007).

65. Ottaviani, C. et al. CD56brightCD16(-) NK cells accumulate in psoriatic skin in response to CXCL10 and CCL5 and exacerbate skin inflammation. eng. Eur J Immunol 36, 118–128. ISSN: 0014-2980 (Print); 0014-2980 (Linking) (Jan. 2006).

66. McKinney, E. F. et al. A CD8(+) NK cell transcriptomic signature associated with clinical outcome in relapsing remitting multiple sclerosis. eng. Nat Commun 12, 635. ISSN: 2041-1723 (Electronic); 2041-1723 (Linking) (Jan. 2021).

67. Fehrmann, R. S. N. et al. Trans-eQTLs reveal that independent genetic variants associated with a complex phenotype converge on intermediate genes, with a major role for the HLA. PLoS Genet 7, e1002197 (Aug. 2011).

68. Norman, P. J. et al. Defining KIR and HLA Class I Genotypes at Highest Resolution via High-Throughput Sequencing. eng. Am J Hum Genet 99, 375–391. ISSN: 1537-6605 (Electronic); 0002-9297 (Print); 0002-9297 (Linking) (Aug. 2016).

