## SUPPLEMENTARY FIGURES 1-6 for "Natural Killer cells demonstrate distinct eQTL and transcriptome-wide disease associations, highlighting their role in autoimmunity"

### Natural Killer cells demonstrate distinct eQTL with implications for immune related disease states

March 24, 2021

Figure S1: **The effects of principal component incorporation for *cis* and *trans* eQTL mapping.** Graphs demonstrate the effect of incorporating between 0 and 50 principal components as covariates on eQTL discovery in *cis* (top) and *trans* (bottom) mapping. The number of eQTLs discovered represent approximate passes, and actual numbers differ in the final analysis.

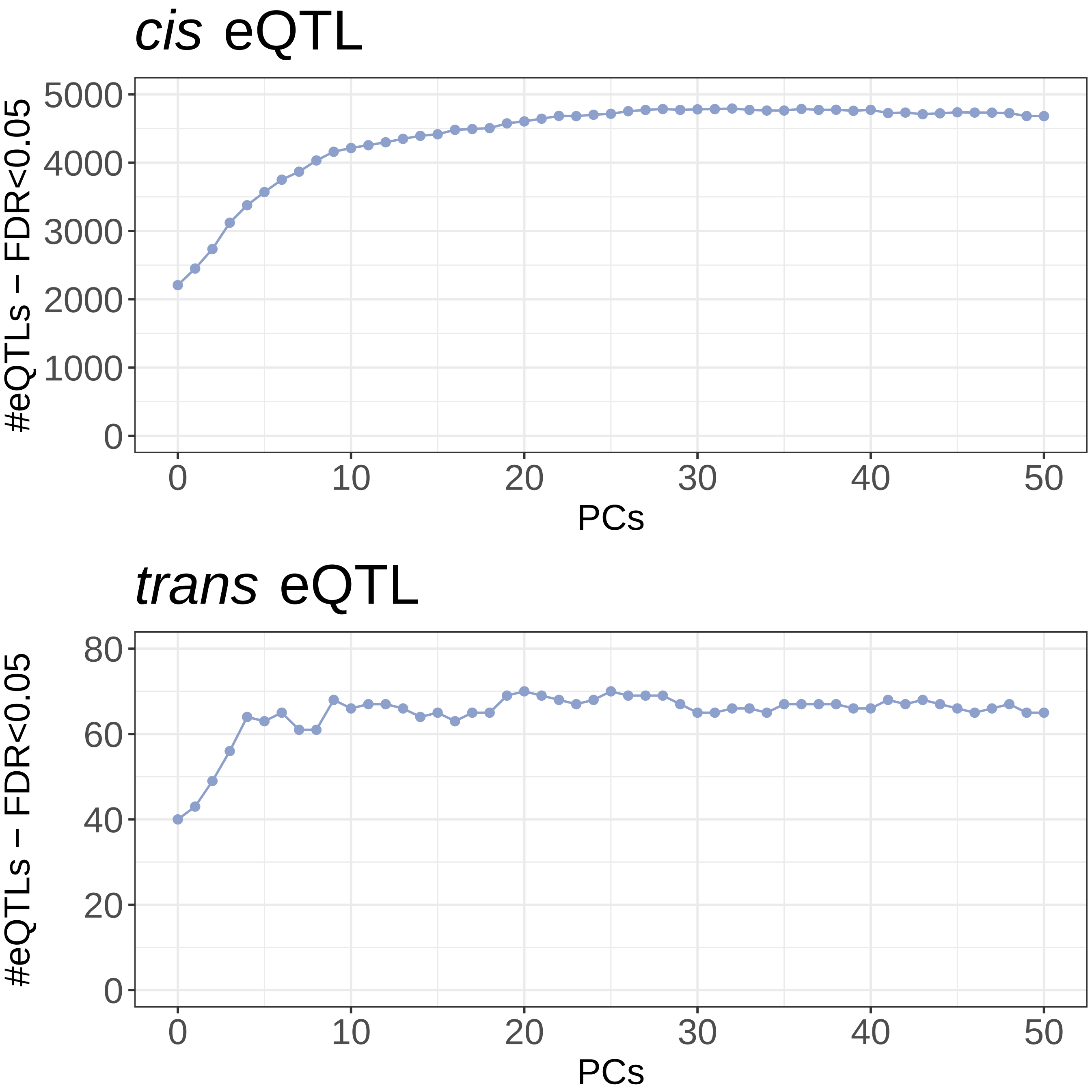

Figure S2: **Functional annotation of NK cell eQTL.**
Frequency plots representing the density of eQTL loci surrounding functional genomic tracks. P-values (permutation) represent enrichment; significantly overrepresented features are coloured pink, underrepresented features blue. Densities are plotted for primary eQTL (A), secondary eQTL (B) and NK cell-specific eQTL (C). Enrichment of eQTL loci at transcription factor binding sites for primary eQTL (D), secondary eQTL (E) and NK cell-specific eQTL (F). Significantly enriched transcription factor binding sites are highlighted (pink). (G) Comparison of functional track and transcription factor binding sites overlap observed in primary and secondary eQTL. P-values calculated with Fisher Exact tests. (H) Comparison of functional track and transcription factor binding sites overlap observed in eQTL specific to NK cells eQTL shared with other cell types. Significantly overrepresented features are coloured pink, underrepresented features blue.

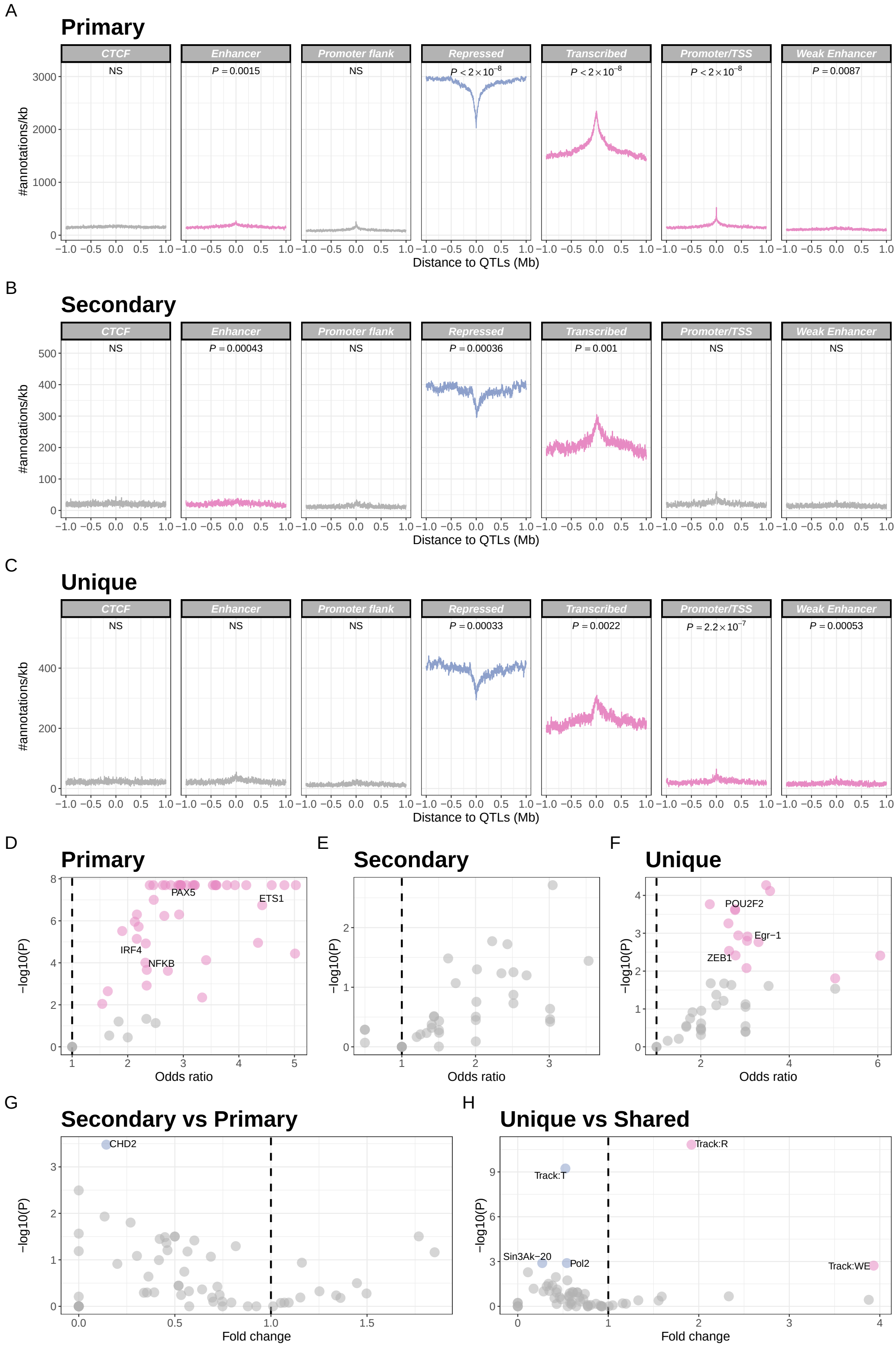

Figure S3: **GOBP enrichment of NK cell-specific eQTL.**
GOBP pathway analysis of genes with eQTL specific to NK cells (as compared with monocytes, neutrophils, $CD4^{+}$ and $CD8^{+}$ T cells. All depicted pathways are significantly enriched ($FDR < 0.05$). nOverlap, number of overlapping genes from a pathway. Enrichment is calculated using a hypergeometric test.

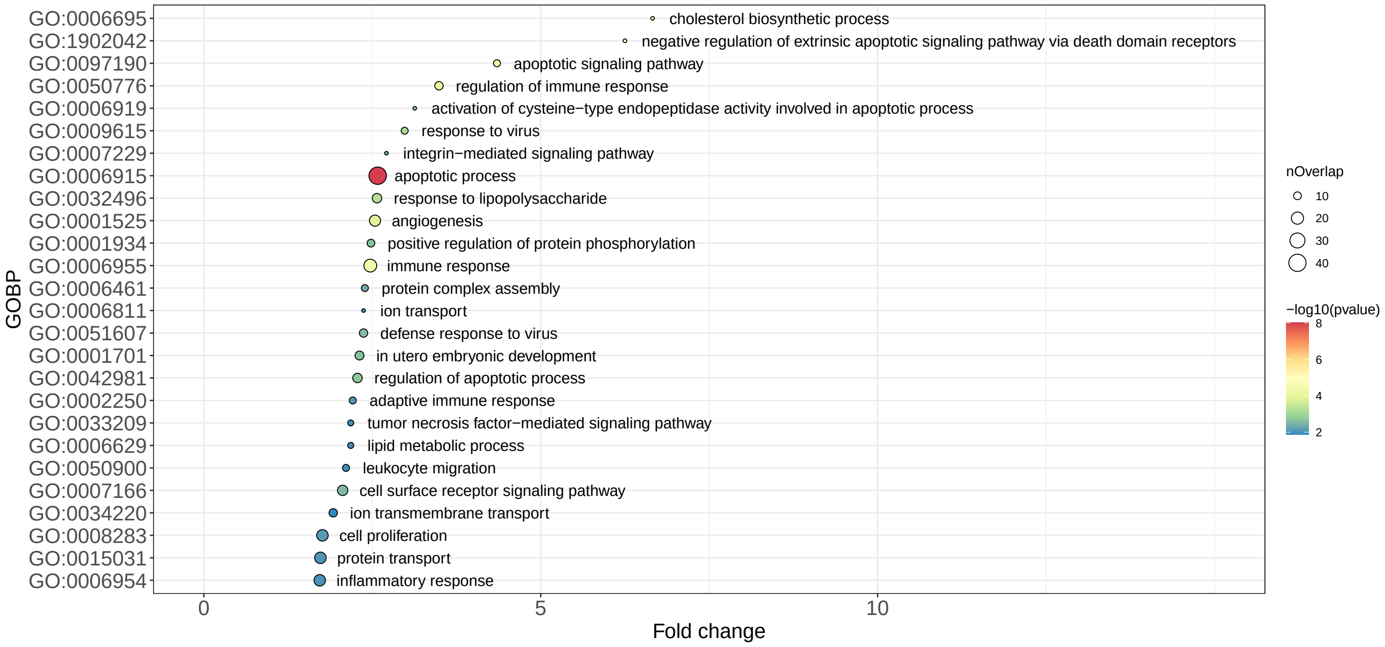

Figure S4: **NK cell eQTL at *ERAP2*.**
(A) Effect of rs1363974 genotype on *ERAP2* expression in NK cells. (B) The effect of rs1363974 genotype on *ERAP2* expression is common to NK cells, monocytes, neutrophils, $CD4^+$ and $CD8^+$ T cells. The *ERAP2* eQTL in NK cells colocalises with a genetic locus which determines neutrophil percentage (C) and lymphocyte percentage (D). SNPs are coloured according to strength of LD (CEU population) to the peak eSNP (rs1363974); brown $r^2 > 0.8$, orange $0.5 < r^2 \leq 0.8$, yellow $0.2 < r^2 \leq 0.5$, grey $r^2 leq 0.2$.

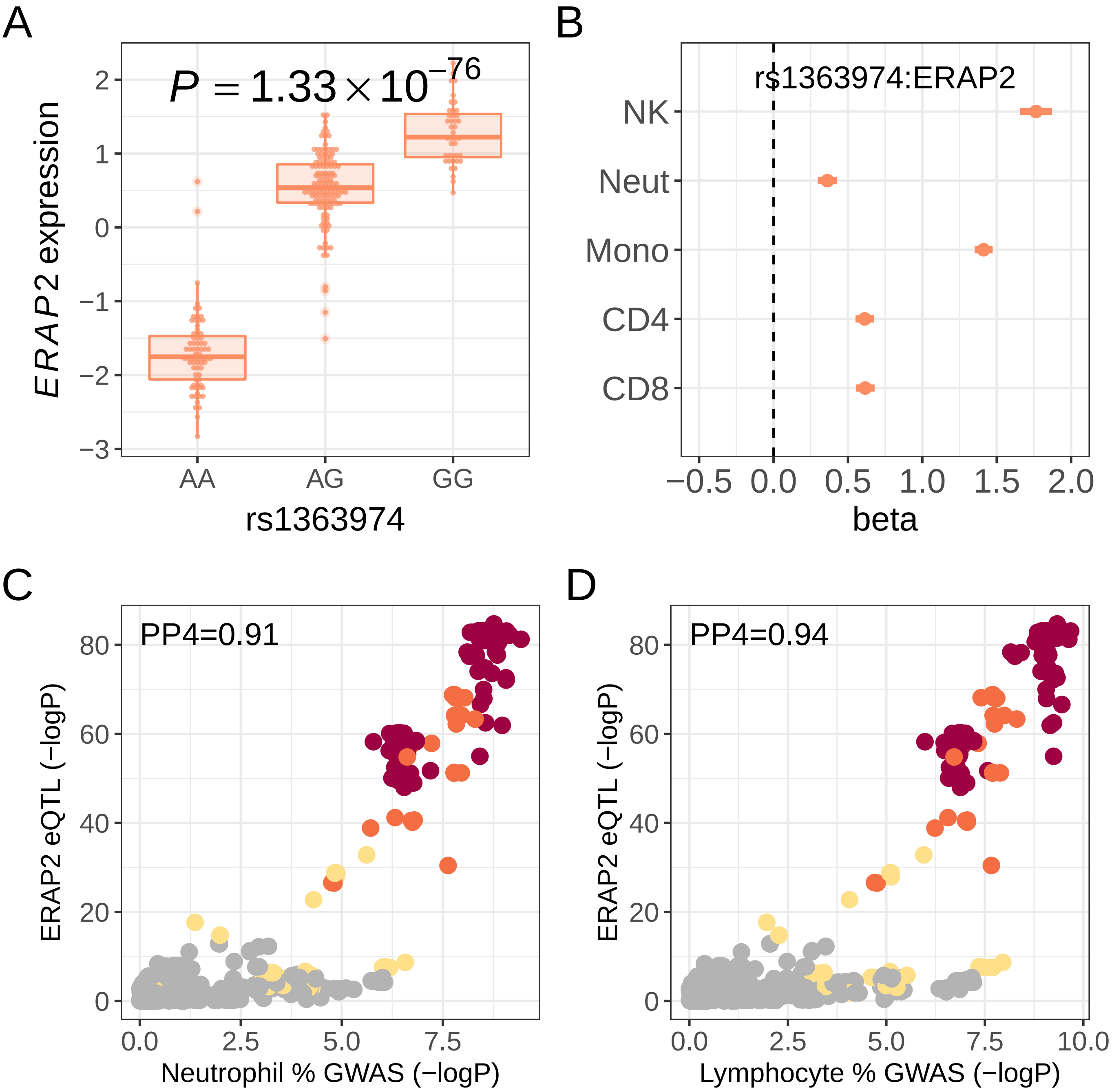

Figure S5:

Allele specific expression was measured at the SNP rs2228479, in $R^2$ of 1 with the peak *cis* eQTL at this gene, rs117406136 across cDNA from individuals heterozygote at this allele using the C-BASE assay in monocytes from 5 individuals, where no difference was seen, and in NK cells. Comparison genomic DNA was used to calculate a $chi^2$ P value.

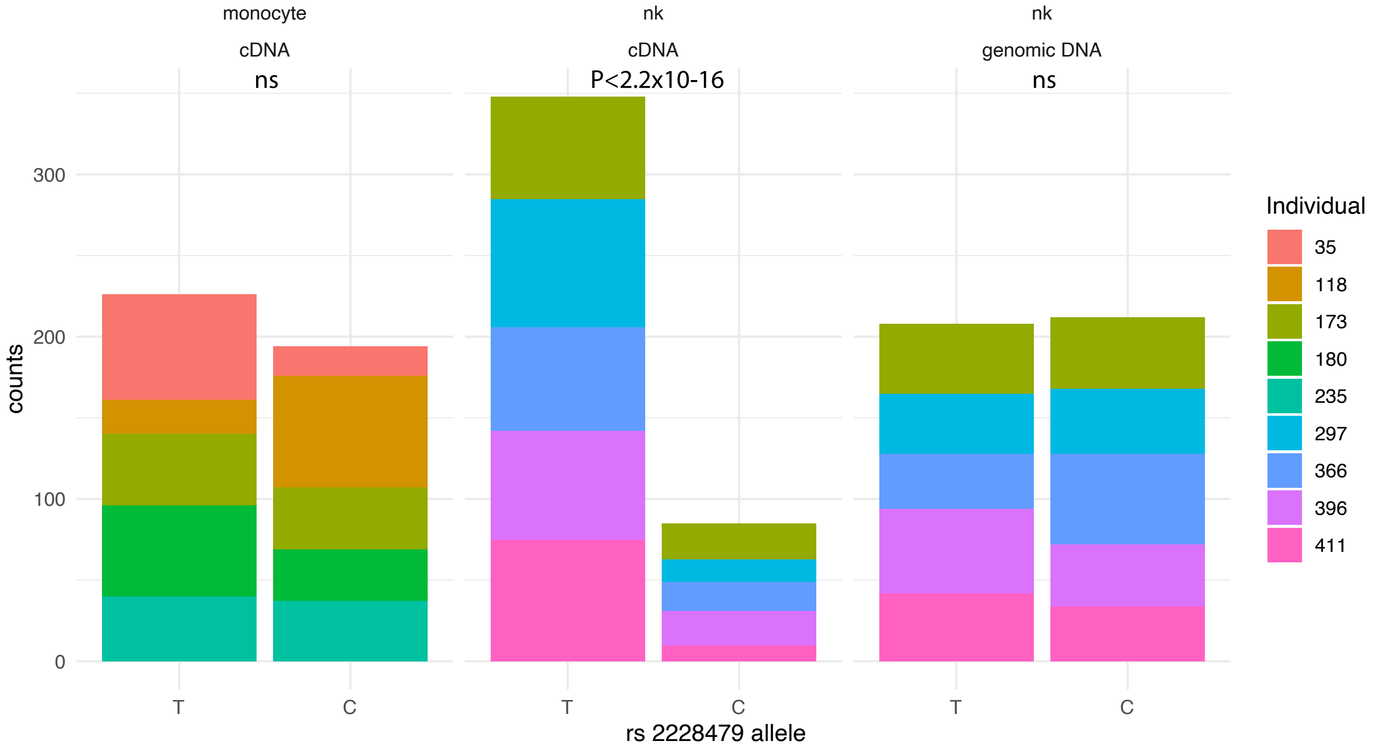

Figure S6: **KIR *trans* eQTL with conditioning on KIR2DS4del.**
Regional association plots depicting effect of KIR region genotype and KIR copy number on gene expression in *trans* for each of 13 genes whose expression is significantly modified by a KIR variant in *trans*. SNPs are coloured grey, KIR types coloured green. Significant associations are highlighted (pink). Associations are plotted unconditionned (left) and conditionned on KIR2DS4del copy number (right).

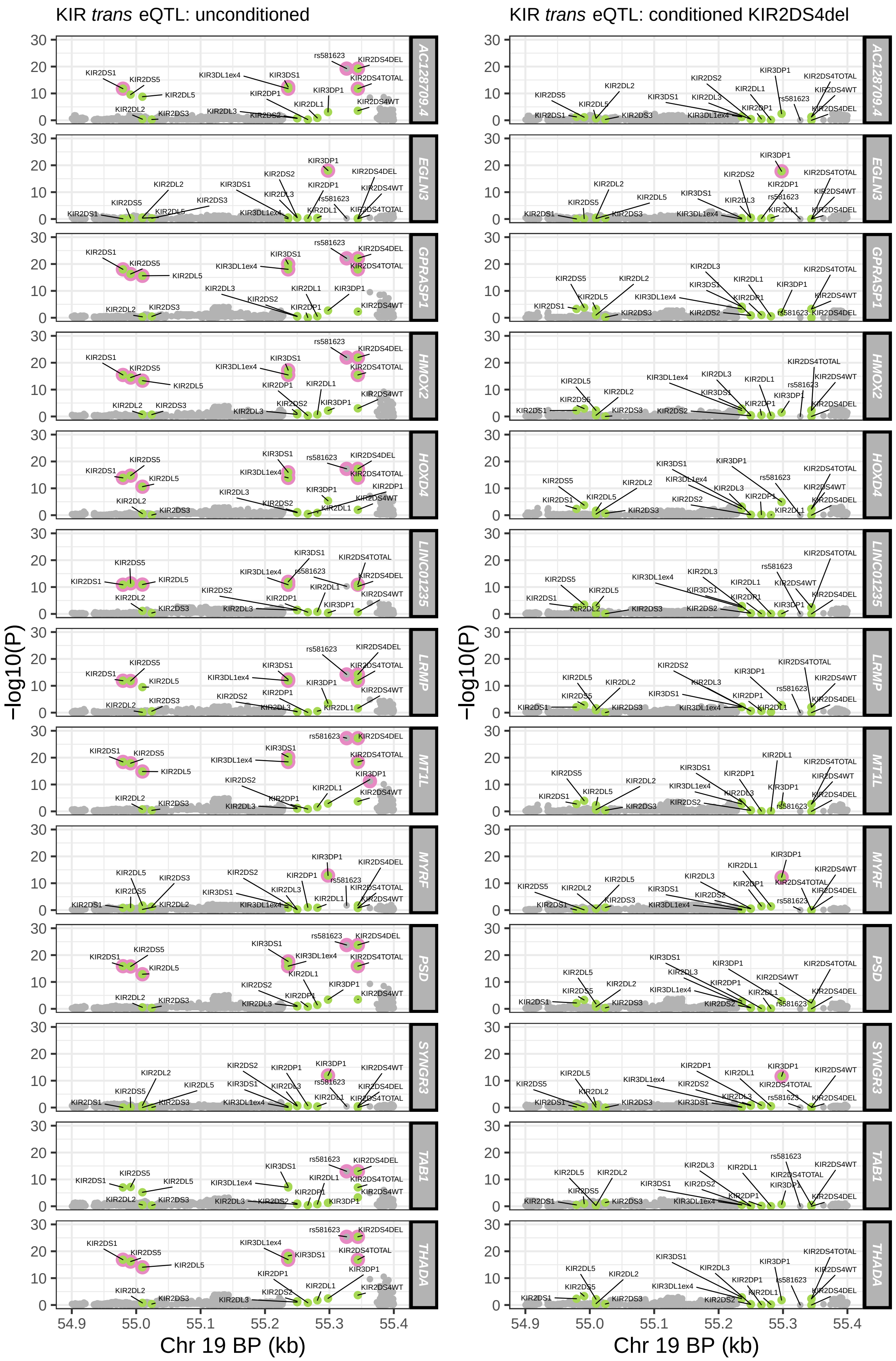
